# Short-term insurance versus long-term bet-hedging strategies as adaptations to variable environments

**DOI:** 10.1101/378091

**Authors:** Thomas Ray Haaland, Jonathan Wright, Jarle Tufto, Irja Ida Ratikainen

## Abstract

Understanding how organisms adapt to environmental variation is a key challenge of biology. Central to this are bet-hedging strategies that maximize geometric mean fitness across generations, either by being conservative or diversifying phenotypes. Theoretical models of bet-hedging and the multiplicative fitness effects of environmental variation across generations have traditionally assumed that environmental conditions are constant within lifetimes. However, behavioral ecology has revealed adaptive responses to additive fitness effects of environmental variation within lifetimes, either through insurance or risk-sensitive strategies. Here we explore whether the effects of adaptive insurance interact with the evolution of bet-hedging by varying the position and skew of fitness functions within and between lifetimes. When insurance causes the optimal phenotype to shift from the peak to down the less steeply decreasing side of the fitness function, then conservative bet-hedging does not generally evolve on top of this, even if diversifying bet-hedging can. Canalization to reduce phenotypic variation within a lifetime is almost always favored, except when the tails of the fitness function are steeply convex and produce a novel risk-sensitive increase in phenotypic variance akin to diversifying bet-hedging. Importantly, using skewed fitness functions, we provide the first example of how conservative and diversifying bet-hedging strategies might coexist.

## Introduction

How organisms adapt to unpredictable fluctuations in the environment has been an intriguing and important problem for many years in evolutionary biology, and especially recently when predicting adaptive responses to environmental change. Conditions may vary over different time scales, selecting for adaptations that maximize fitness in the face of environmental stochasticity in everything from labile behavioral traits within a lifetime (e.g. variance-sensitive foraging, Stephens 1981) to cross-generational effects of life-history traits (e.g. bet-hedging; Simons 2011; Starrfelt and Kokko 2012). Thus, the phenotypes we observe in organisms today have likely been shaped by environmental variation experienced across longer timescales during their evolutionary history, and trait values may not necessarily appear optimal when considering just short-term current environmental conditions (Nadeau et al. 2017). Environmental variation itself is expected to be a strong selective agent, since genotypic rather than individual fitness determines optimal strategies that are produced over evolutionary time in stochastic environments (Lewontin and Cohen 1969; McNamara 1998).

Asymmetric fitness functions pose an additional challenge to evolutionary biologists seeking to understand genotypic adaptations in variable environments (Yoshimura and Shields 1987; Urban et al. 2013). Skew in the function relating a single, continuous phenotypic trait to fitness is commonly seen in nature, occurring whenever costs and benefits differ in how they relate to increasing versus decreasing values of the phenotype, or when the strength of selection acting on the two sides of the phenotypic distribution differs. Common examples are thermal performance curves (Angilletta 2009), optimal clutch or litter sizes (Mountford 1968; Boyce and Perrins 1987), and reproductive benefits versus viability costs of sexually selected ornaments (Andersson and Iwasa 1996). In these types of scenarios, uncertainty across instances in any component determining individual fitness will cause the optimal trait value to differ from the trait value at the peak of the fitness function (Yoshimura and Shields 1987; Parker and Smith 1990). Such uncertainty in fitness pay-offs across instances is also almost ubiquitous in biological systems. Across lifetimes, phenotypic differences among individuals (as instances) of the same genotype may arise due to developmental instability creating random (uncanalized) variation in phenotypes and thus also in their fitness, and avoiding such variation may be costly (Zhang and Hill 2005). Within lifetimes, uncertainty may occur in individual energetic state on short (e.g. behavioral) timescales, due to stochastic probabilities of resource acquisition such as prey captures. In addition, the fitness effects of the phenotype itself (i.e. the shape or position of the fitness function) may be uncertain, for example due to micro-environmental variability, or variation occurring over short time scales, such as in social environments.

With a skewed fitness function, any stochastic environmentally induced variation in fitness pay-offs will select for apparently suboptimal phenotypes with trait values away from the peak of the deterministic fitness function when selection maximizes arithmetic mean fitness (Fig. 1). Finding the (arithmetic) mean fitness in such cases involves multiplying the phenotype-specific fitnesses with the frequencies of the different phenotypes (Mountford 1968). This phenomenon is sometimes described as the cliff-edge effect (Vercken et al. 2012; Mitteroecker et al. 2016), and is commonly encountered as ‘insurance’ strategies in fields such as behavioral ecology (Dall 2010). A well-known example is the small bird in winter (Brodin 2007). Facing a starvation-predation trade-off, the small passerine bird wants to be as light as possible to nimbly avoid predators during the day, but needs to store fat before nightfall, which it metabolizes to stay warm during the night. The small bird in winter will therefore adaptively store more fat when temperatures are more variable (Bednekoff et al. 1994), and/or when food supply is more uncertain (Krams et al. 2010; Ratikainen and Wright 2013)

**Figure 1:**
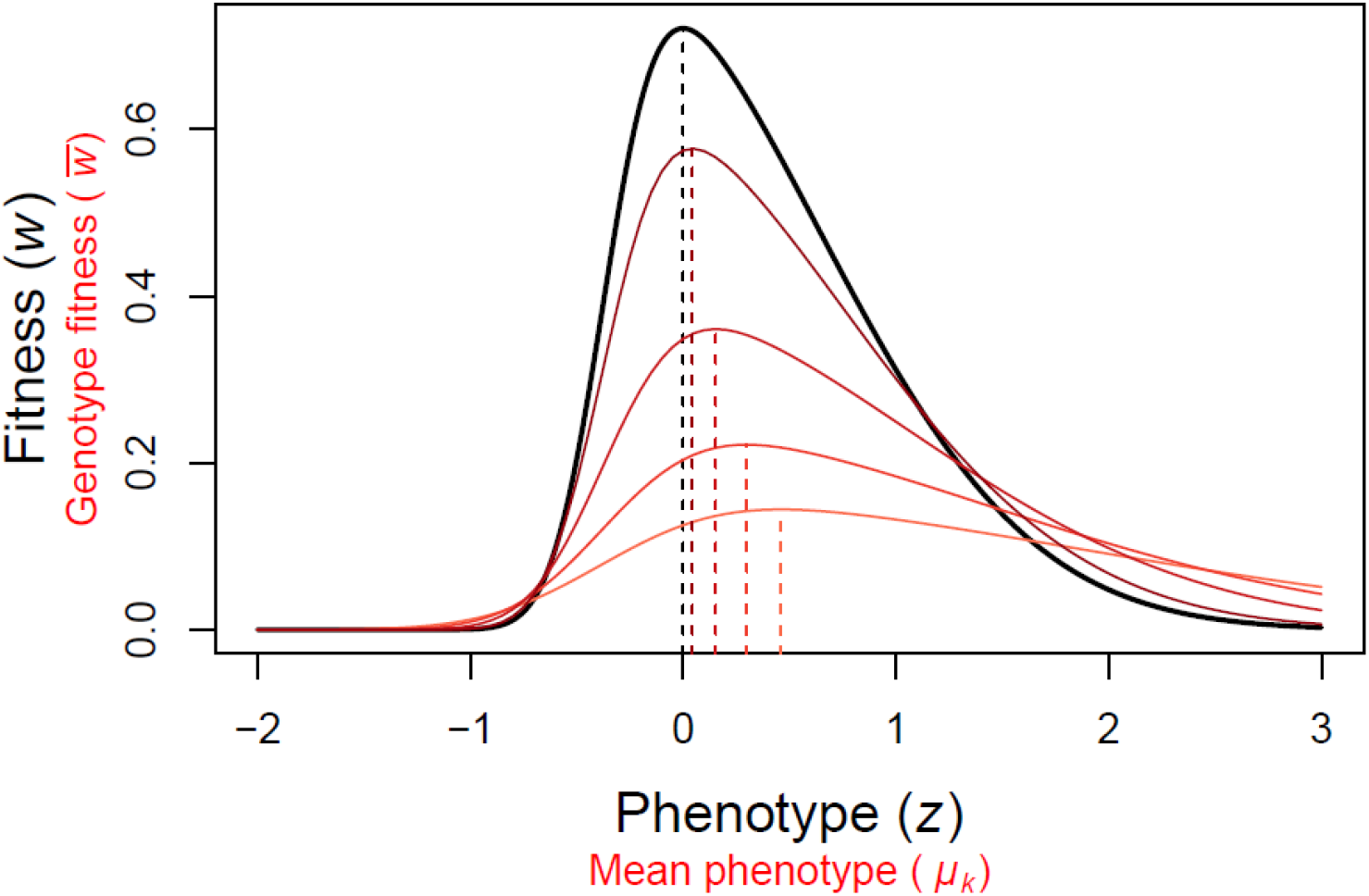
Asymmetric fitness functions and phenotypic variation produce insurance. Black line: the skew normal fitness function with location *θ* = *θ*_0_, width *ω* = 1 and skew *α* = 5. Red lines: arithmetic mean fitness 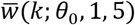 across individuals for genotypes with a fixed *σ_k_*, from darkest to lightest, *σ_k_* = {0.5, 1, 1.5, 2}. Maxima of these functions are indicated by the dotted lines, and approximate to 0.002, 0.044, 0.153, 0.297 and 0.458, respectively.

Across generations, however, the fitness of a lineage is determined by its geometric mean fitness rather than the arithmetic mean, due to reproduction being an inherently multiplicative process (Lewontin and Cohen 1969; Simons 2011). When environmental conditions are constant between generations (i.e. the function relating the trait in question to fitness is exactly the same each generation), arithmetic and geometric mean fitness are equal. However, once some aspect of the fitness function differs between generations, creating variation in realized fitness between individuals with the same genotype, then the geometric mean will be lower than arithmetic mean. Crucially, a change in strategy that lowers variance in realized fitness at the genotype level may increase geometric mean fitness, and thus be selectively favored. If such a strategy that increases geometric mean fitness at the genotype level also involves a simultaneous decrease in arithmetic mean fitness, it is defined as bet-hedging (Philippi and Seger 1989). Two main types of bet-hedging are usually considered, diversifying bet-hedging (DBH) and conservative bet-hedging (CBH) (Philippi and Seger 1989; Simons 2011; Starrfelt and Kokko 2012). DBH increases phenotypic variance and thus reduces the correlations in fitness between individuals of the same genotype, such that not all individuals are affected by the environment in the same correlated way. DBH strategies can include both producing two discrete ‘types’ of individuals, such as dry and wet-year specialists, or dormant and active life stages in response to good and bad years (Lewontin and Cohen 1969; Venable 2007; Graham et al. 2014), and continuous variation in a trait among individuals, such as size of offspring, or timing of reproduction and entering or exiting dormancy (Simons 2009; Devaux and Lande 2010; Lof et al. 2012). In contrast, CBH reduces variance in realised fitness at the individual level, such that each individual within a genotype will perform moderately well across a range of environments. However, there is nothing stopping a strategy from combining the two types, and Starrfelt and Kokko (2012) argue that DBH, reducing only among-individual fitness variance, and CBH, reducing only each individual’s fitness variance, are actually two ends of a continuum with strategic combination of DBH and CBH possible in between. Despite this, neither Starrfelt and Kokko (2012) nor later authors exploring similar models (e.g. Crowley et al. 2016) explain how such a combination of DBH and CBH would work. These papers examine models with two discrete environments, and a suggested conservative bet-hedger (acting as a generalist coping moderately well with both environments) is never able to outperform a diversified bet-hedger (producing specialists to each environment in the proportions that they occur).

We present a different interpretation of CBH, which potentially allows for both CBH and DBH to coexist within the same model. Considering a continuous trait with an asymmetric fitness function that fluctuates between generations, CBH can be envisioned as having a cliff-edge effect in the same way as insurance (see above). Organisms would thus be ‘playing it safe’ by shifting the mean trait value away from the fitness function maximum, towards the less steeply decreasing side. In such a scenario, we expect insurance to maximize arithmetic mean fitness within a generation. An additional shift in the optimal trait value even further away from the cliff edge might then be selected for if it lowers fitness variance between generations (despite lowering arithmetic mean fitness in a single generation). Such an effect would essentially constitute a CBH strategy. Phenological features such as breeding date, migration date or egg laying date for temperate birds are examples of traits with such asymmetric fitness functions. The strength of selection may differ for the underlying selection pressures, for example if being too late leads to lower offspring competitive ability, but being too early leads to a much more severe mismatch with the food peak resulting in complete reproductive failure (Gienapp 2012). Whether breeding after the peak in the fitness function represents insurance or CBH depends upon whether the mismatch between the mean trait value and the peak of the asymmetric fitness function is the result of individuals maximizing arithmetic mean fitness within their lifetime versus lineages being favored that maximize geometric mean fitness over long time periods (see Lof et al. 2012). Despite having much in common and some confusion between the terms in the literature, CBH has rarely been placed in the same theoretical framework as insurance, and insurance has been all but absent as part of the bet-hedging literature.

Here we investigate the relative importance of insurance and CBH in coping with stochastically fluctuating environments within and between generations when the fitness function is skewed. We use a single, continuous trait and calculate the means and variances in phenotype that maximize arithmetic or geometric mean fitness under different magnitudes of fluctuations in the optimal trait value. Mechanisms regulating the phenotypic variance expressed within a genotype, such as DBH increasing such variance or phenotypic canalization decreasing it, are expected to interact with insurance and/or CBH. Previous theoretical work has shown that DBH will adaptively increase variance in trait values once the variance in the phenotypic optimum exceeds the squared width of the fitness function, whereas smaller environmental variance favors the opposite mechanism, canalization of the trait towards the value that maximizes fitness in the mean environment (Slatkin and Lande 1976; Bull 1987). Intuitively, greater stochastic variation in trait values should require there to be more insurance or CBH modifying the mean trait value, but these different components have not previously been placed in a common framework. By using a skewed fitness function to illustrate the effects of insurance versus CBH, we are able to examine these interactions, whilst modeling DBH alongside CBH in such a way allows us to formally explore Starrfelt and Kokko’s (2012) suggestion regarding an adaptive continuum between these two potentially coexisting forms of bet-hedging.

## Model description

### The skew normal fitness function

A wide variety of fitness functions have been used to characterize asymmetrical relationships between phenotype and fitness (Martin and Huey 2008; Vasseur et al. 2014), but the results we demonstrate here can arise from any function with nonzero skew. We base our skewed fitness function on the density function of the skew normal distribution (O’Hagan and Leonard 1976), omitting the normalizing constant such that individual fitness takes a value of one for *z* = *θ*. Fitness as a function of the phenotype *z* is then given by

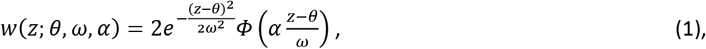

which is a Gaussian function multiplied by a term involving the cumulative distribution function *Φ* of the standard normal distribution. The parameter *θ* specifies the location, *ω* the width, and *α* the skew of the fitness function. *α* > 0 gives a right-skewed function (positive skewness), *α* < 0 a left-skewed function, and *α* = 0 a symmetric Gaussian fitness function. Importantly, *α* also changes the position of the maximum value of *w* (despite location parameter *θ* being kept constant), so we will write *θ*_0_ as the value of *θ* that provides maximum fitness for a trait value of zero. To examine the effects of skewed fitness functions on trait values we will use *α* > 0 and *θ* = *θ*_0_, so that adaptations in terms of phenotypic values shifted away from the fitness function maximum (due to insurance or CBH) become positive and easily interpretable relative to zero (i.e. the value of *z* simply becomes the distance from the peak, or the ‘amount’ of insurance or CBH).

### Genotypic fitness when phenotypes vary within genotype

There may be variation in the phenotype *z* produced by a certain genotype *k*, due either to some (adaptive or non-adaptive) instability in individual development, or due to different individuals experiencing different microclimates. Following Bull (1987), we assume this variation to follow a normal distribution *f_k_*(*z*), with a mean *μ_k_* and variance *σ*^2^_*k*_. We are interested in the joint evolution of the two underlying genotypic values *μ_k_* and *σ_k_*, and assume no genetic linkage or pleiotropic effects between them.

The mean fitness of all individuals with the genotype *k* (with genotypic values *μ_k_* and *σ*^2^*k*) in any given environment *θ* (or a constant environment over time) then becomes

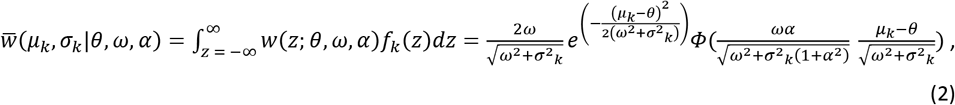

which is akin to eq. (3) in Bull (1987). The resulting function 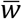 has the same form as (1) but with a larger width parameter and a smaller skew parameter. When *σ*^2^*_k_* = 0, the functions are identical. Choosing a constant phenotypic variance *σ^2^_k_* > 0, we can use numerical optimization over *μ_k_* and compare the difference in maxima of *w* and 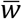, to find the amount of insurance needed to maximize arithmetic mean fitness across all individuals of the genotype (i.e. the optimal shift in mean phenotype away from the fitness function maximum). Fig. 1 shows *w* (black) together with 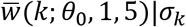 (reds) for increasing values of *σ_k_*. In this case, since the maximum of the fitness function is at zero, the optimal amount of insurance is simply the value of the phenotype that gives the highest fitness, 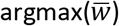. The larger the phenotypic variance *σ*^2^*k*, the more insurance is needed to maximize genotype fitness. We also note that genotype fitness strongly declines with increasing phenotypic variance - except when the mean phenotype is far away from the fitness function peak.

### Long-term fitness in a fluctuating environment

In a fluctuating environment it is not just individual fitness but the fitness of a genotype that will differ in different environments, and long-term fitness in such cases is determined by geometric mean rather than arithmetic mean fitness (Lewontin and Cohen 1969; Simons 2011). In the case of no fluctuations, the geometric mean is simply equal to the arithmetic mean, and equation (2) is valid. If we let the optimum position *θ* follow a normal distribution *f* with a mean of *θ*_0_ and a variance *σ_θ_* across generations, long-term arithmetic mean fitness can be found by taking the expectation of equation (2) across the environmental fluctuations,

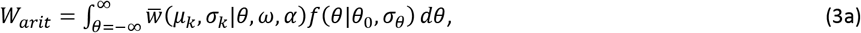

and for geometric mean fitness we can take the exponential of log fitnesses integrated across all different environmental conditions,

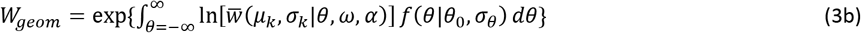

Maximizing both eq. 3a and 3b over *μ_k_* and *σ_k_* gives the strategy that provides the highest long-term arithmetic or geometric mean fitness, respectively, for genotype *k*. Comparing them allows us to tease apart bet-hedging effects (those maximizing geometric mean fitness at the expense of arithmetic mean fitness) from non-bet-hedging effects (those maximizing only arithmetic mean fitness). Maximizing 3a for a fixed *σ_k_* gives the optimal amount of insurance (since the peak of the individual fitness function is at zero), equal to argmax_*μ*_ *W_arit_*|*σ_k_*. Maximizing 3b for the same *σ_k_* then reveals whether any additional shift in mean phenotype can be attributed to bet-hedging rather than just insurance, with the optimal amount of CBH then being equal to argmax_*μ*_[*W_arit_*|*σ_k_*] - argmax_*μ*_ [*W_arit_*|*σ_k_*].

Numerical integration was carried out in R version 3.3.1 (R Core Team 2016) and the code is available upon request.

## Results

Figures 2 and 3 show the long-term fitness of genotype *k* (consisting of the gene for mean phenotype, *μ_k_*, and for variance in the phenotype, *σ^2^_k_*) measured in either arithmetic or geometric mean fitness in an environment with increasing fluctuations in the position parameter *θ* of the individual fitness function (increasing *σ_θ_*). In Fig. 2 the individual fitness function is symmetrical and in Fig. 3 it is skewed. With a stable environment across generations (*σ_θ_*=0) the arithmetic and geometric mean fitness are equal (top and bottom panels the same). The fitness surface in these cases peaks at *μ_k_*=0 and *σ*^2^_k_=0, that is, the
 optimal genotype is a trait value phenotypically canalized (i.e. with as little variation as possible) at the peak of the individual fitness function. As environmental fluctuations increase (*σ_θ_*>0), the differences between arithmetic and geometric mean fitness increase. Notably, in Fig. 2 the peak geometric mean fitness contours move upwards in the bottom panels as the environmental fluctuations increase. This adaptive increase in phenotypic variation (*σ_k_*) within the genotype represents DBH, and since arithmetic mean fitness strictly declines with increasing *σ_k_* (top panels), the necessary requirement that bet-hedging involves a lowering of arithmetic mean fitness is fulfilled. In accordance with Bull’s (1987) result, this selection for increased phenotypic variation only appears once the environmental variance *σ*^2^_*θ*_ is larger than the squared width of the fitness function, and the optimal *σ*^2^_*k*_ is then equal to *σ*^2^_*θ*_ - *ω*^2^. For the symmetric Gaussian distribution (Fig. 2) this threshold is simply *ω*^2^ = 1, and DBH appears when *σ*^2^_*θ*_ > 1. In Fig. 3 asymmetry is introduced into the fitness function (results are shown for *α* = 5, which matches the fitness function in Fig. 1), but all other parameters remain as in Fig. 2. The variance of the fitness function decreases as the skew increases, so this scenario also produces DBH (fitness surface peak with *σ*^2^_*k*_ > 0) under lower levels of environmental variance. The skew also leads to the fitness surface peaks shifting to positive values of *μ_k_*, away from the steeply decreasing side of the individual fitness function. This shift is seen both in the top and bottom rows of Fig. 3, and in each case the shift occurs to the same extent for the maximization of arithmetic and geometric mean fitness. Therefore, this represents only insurance and there is no extra shift that can be attributed to a CBH effect if *μ_k_* and *σ_k_* are jointly maximized.

**Figure 2:**
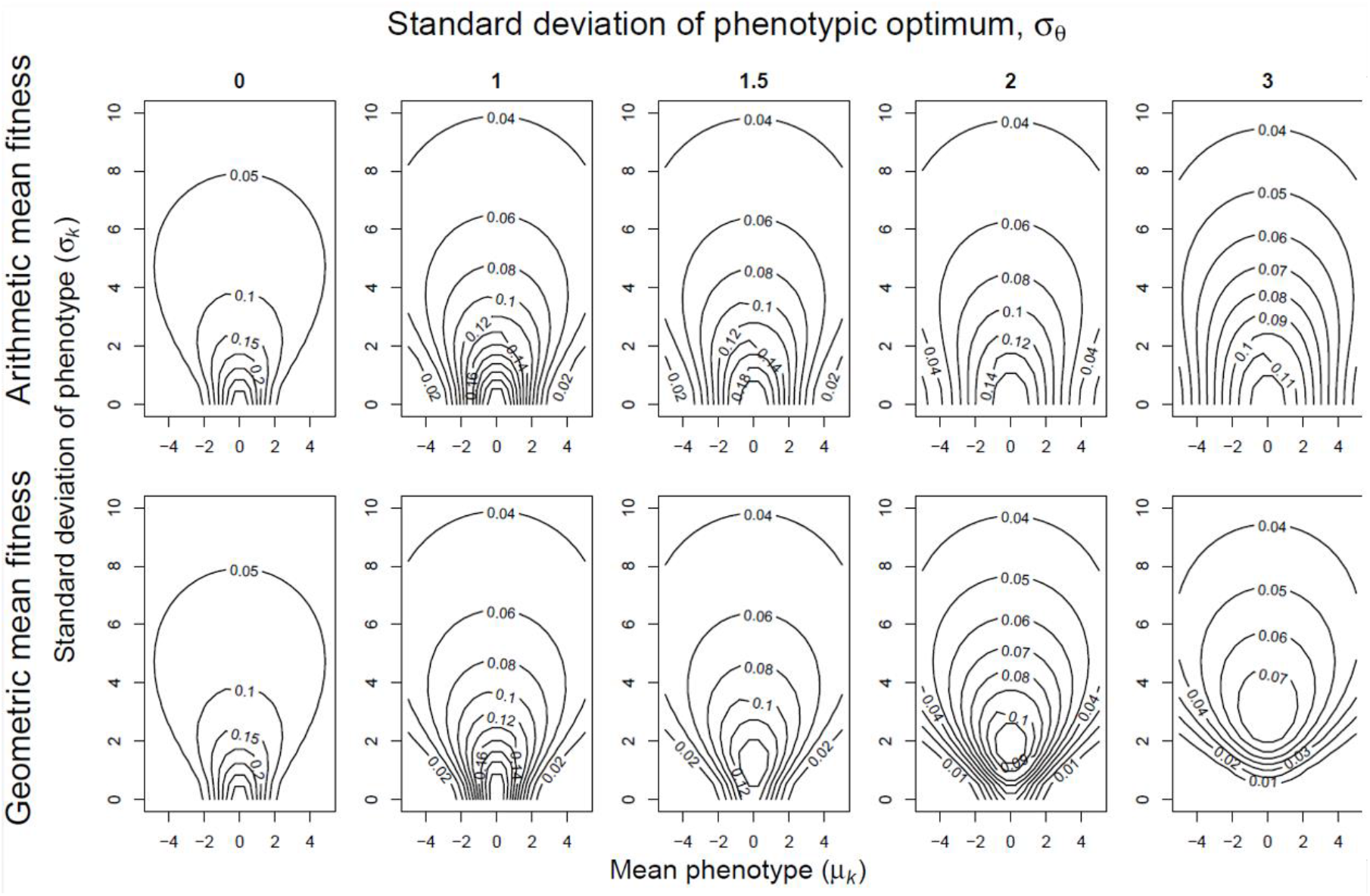
Fitness surfaces for a symmetrical fitness function for genotypes with different values of the mean phenotype *μ_k_* and variation in phenotype *σ_k_*. Contour lines show long-term arithmetic mean fitness (top row) and geometric mean fitness (bottom row). The position *θ* of the fitness function fluctuates stochastically between generations, *θ*^~^N(0, *σ_θ_*), the magnitude *σθ* of environmental fluctuations increases successively (from 0 to 3) in the different columns from left to right. Irrespective of the scale of these fluctuations and the phenotypic variation (*σ_k_*), fitness is always maximized by a peak in the contours in the middle of the x-axis, corresponding to a mean phenotypic value (*μ_k_*) of zero, because the individual fitness function is a symmetrical normal distribution with a mean of 0 and width of *ω*=1.

**Figure 3:**
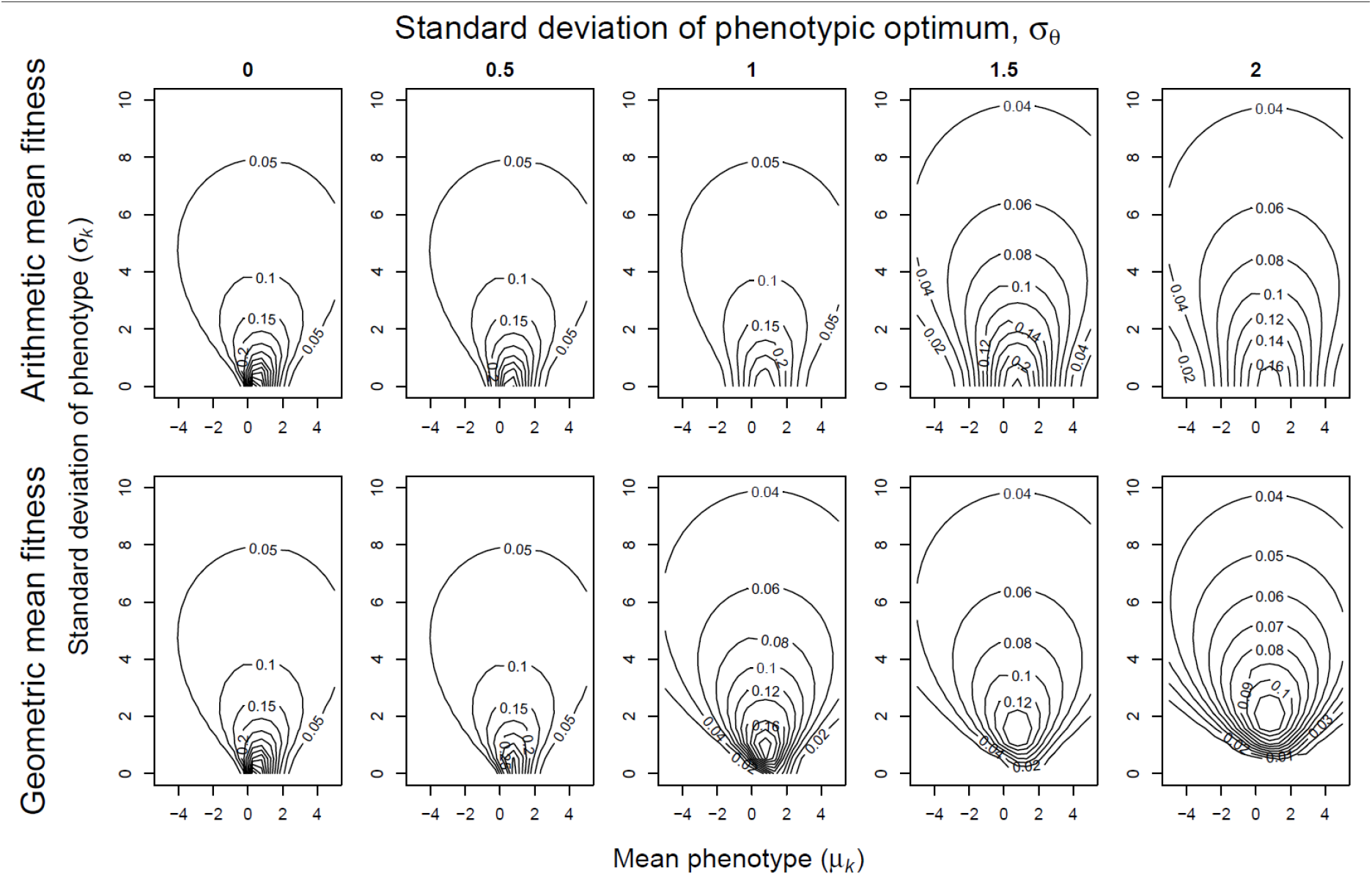
Fitness surfaces for an asymmetrical fitness function for genotypes with different values of the mean phenotype *μ_k_* and phenotypic variation *σ_k_*. Contour lines show long-term arithmetic mean fitness (top row) and geometric mean fitness (bottom row). The position *θ* of the fitness function fluctuates stochastically between generations, *θ*^~^N(0, *σ_θ_*), the magnitude *σ_θ_* of the fluctuations increasing from left to right. In contrast to Fig. 2, fitness is always maximized here by a peak in the contours to the right of the middle of the x-axis (where *μ_k_*=0), because the individual fitness function is a skew normal distribution with a mode of 0, width of *ω*=1 and skew of *α*=5.

However, geometric mean fitness is maximized for higher mean trait values *μ_k_* than arithmetic mean fitness when phenotypic variation *σ_k_* is constrained at low values - see Figs 4 & 5a. Fig. 4 shows this difference in mean phenotype (argmax_*μ*_ *W_geom_* - argmax_*μ*_ *W_arit_*), which is attributable to CBH (y-axis), for different amounts of phenotypic variation *σ_k_* (x-axis) and environmental variation *σ_θ_* (line color). Note that this CBH effect is only detectable when phenotypic variation *σ_k_* is small, and environmental fluctuations *σ_θ_* are large enough that DBH would provide a much greater fitness gain (the steepest incline on the fitness surface comes by moving upwards along the *σ_k_* axis). The selection gradient will thus adaptively increase the phenotypic variance, and not the mean phenotype *per se*.

**Figure 4:**
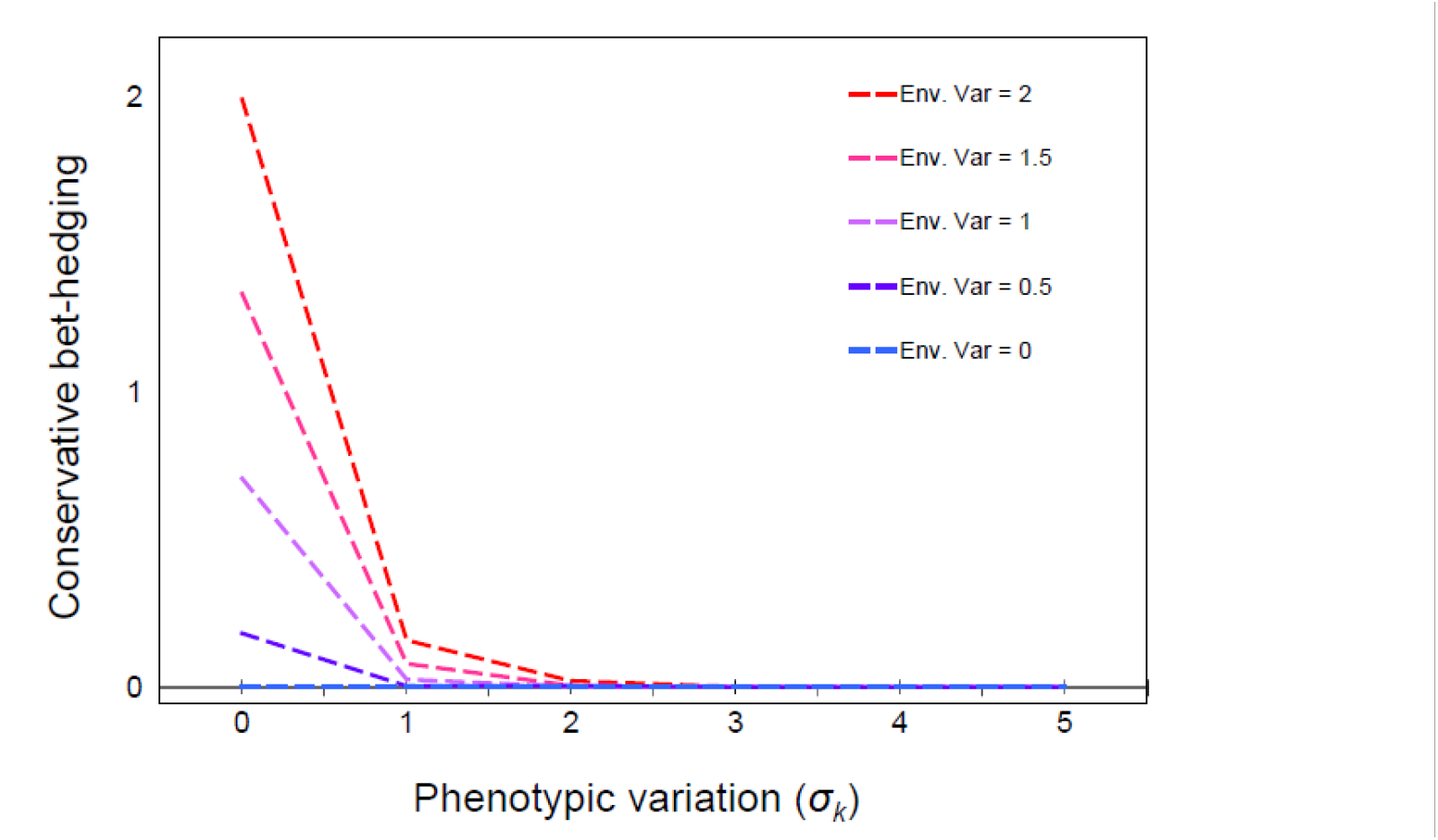
Conservative bet-hedging (CBH), defined as the difference in the mean phenotype (*μ_k_*) maximizing long-term arithmetic versus geometric mean genotype fitness, for given phenotypic standard deviation (*σ_k_*). Lines represent results for different values environmental variation *σ_θ_*, as in Fig. 3. The individual fitness function *w* has a width of *ω*=1 and skew of *α*=5, also as in Fig. 3.

For phenotypically canalized traits (i.e. traits that have experienced selection in *σ_k_* towards zero), we can infer that the fitness functions of these traits have fluctuated less across generations (smaller *σ_θ_*), as we otherwise would not see have seen this canalization. For these values of *σ_θ_*, geometric and arithmetic mean fitness peaks at more similar *μ_k_* values when *σ_k_* = 0 (fig. 4, bluer lines). As *σ_k_* increases, mean fitness 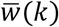 becomes less skewed (see eq. 2, skew parameter *α* decreases with increasing *σ_k_*) and therefore CBH will not shift *μ_k_* on top of any insurance already occurring. We note that if *σ_k_* is constrained to only exhibit a limited amount of DBH, then CBH and DBH will co-occur (e.g. the dependence of optimal *μ* on *σ* in the lower right panel of fig. 3; redder lines showing CBH > 0 at *σ_k_* = 1 in fig. 4), but there are no cases in Figs. 3 and 4 where CBH co-occurs with optimal amounts of DBH. Once environmental fluctuations have caused DBH (*σ_θ_* > *ω*) and phenotypic variation *σ_k_* increases to its new optimum, geometric and arithmetic mean are maximized at the same mean phenotype, and the shift in mean phenotype relative to the fitness function peak is then attributed to insurance rather than CBH.

Figure 5b illustrates a similar case of an apparent bet-hedging effect being instead attributable to simply maximizing arithmetic mean fitness, and thus not necessarily representing bet-hedging at all. In this case, if the mean phenotype is constrained at a suboptimal value, such as may be the case in a climate change scenario shifting the position of the fitness function, a positive amount of phenotypic variance is adaptive (i.e. a risk-prone strategy due to risk sensitivity (Caraco et al. 1980; Stephens 1981), later termed variance sensitivity - see Discussion). This result can be understood, like DBH, as the different individuals of the genotype being phenotypically different (so that at least some are well adapted to the current conditions) rather than everyone being possibly maladapted. However, there is no need here to invoke a geometric mean (bet-hedging) argument because this diversification of phenotypes simply maximizes arithmetic mean fitness across the individuals of the genotype. This type of adaptive phenotypic variance is often attributed to bet-hedging without considering whether the trait specifically increases geometric mean fitness at the cost of a decrease in arithmetic mean fitness (Mountford 1968).

**Figure 5:**
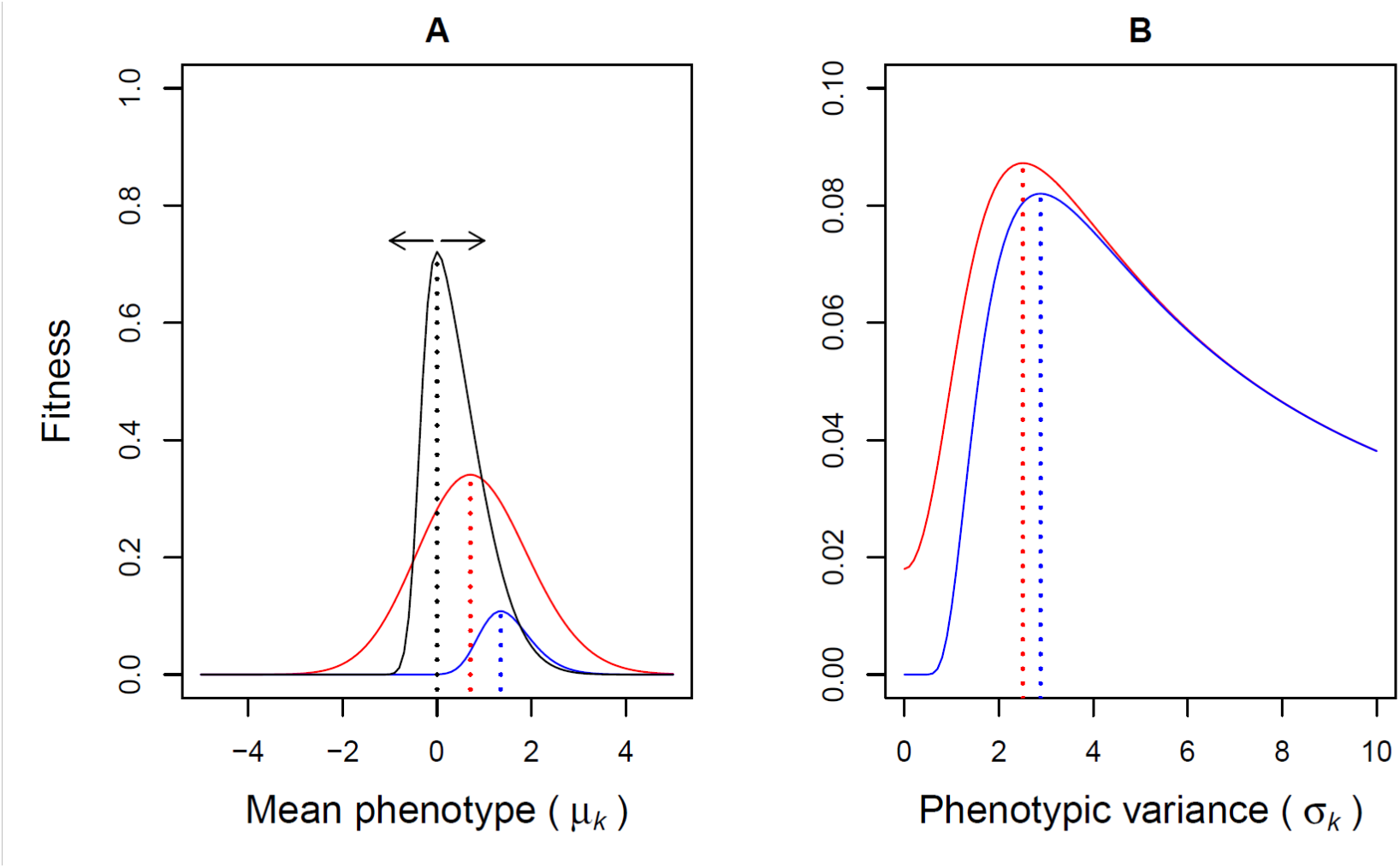
Cross-sections of fitness surfaces in the third column of Fig. 3 (*σ_θ_*=1). The red line is the fitness surface for arithmetic mean fitness, the blue line is the fitness surface for geometric mean fitness. Dotted vertical lines show the maxima for the respective functions. A: a horizontal cross-section taken in the trait (*μ_k_*) dimension at *σ_k_* = 0, hence the individual fitness function (black line) peaks at zero, but fluctuates over time (black arrows), which causes mean fitness to be maximized at positive values of mean phenotype for both arithmetic and geometric mean. B: a vertical cross-section taken in the phenotypic variance (*σ_k_*) dimension at *μ_k_* = -2, and so diversifying bet-hedging is favored by geometric mean fitness, but arithmetic mean fitness also increases with phenotypic variance, because of Jensen’s inequality (the individual fitness function is strongly convex at *μ_k_* = -2) and hence what is known as a variance-prone strategy – see text for details.

## Discussion

Among the various types of adaptive strategies to cope with environmental stochasticity, many have typically been considered from a within-individual perspective in the tradition of behavioral ecology. These use optimality theory to maximize some fitness proxy using the arithmetic mean across instances within a single generation, such as energy intake per time (Davies et al. 2012). An example is optimal foraging, a large body of the behavioral ecology literature that deals with such within-individual traits, including risk-sensitivity, state-dependent energy budgets, adaptive levels of energy reserves, and information sampling of foraging options (Stephens et al. 2007). Some rather different strategies have been considered to operate among individuals and over many generations. This long-term perspective has been in the tradition of evolutionary theory, which has identified key concepts such as phenotypic canalization and bet-hedging (Slatkin and Lande 1976; Bull 1987; Philippi and Seger 1989; Frank and Slatkin 1990). In an attempt to reconcile these contrasting views, we have calculated both long-term arithmetic and geometric mean fitnesses for combinations of trait means and variances under different levels of stochastic environmental fluctuations. By comparing the results obtained when maximizing arithmetic versus geometric mean fitness, we have illustrated some possible similarities between the two approaches, and the discrepancies that arise when considering the effects of either of these two measures of fitness in isolation.

Crucially, we use skewed fitness functions to demonstrate that shifting the mean phenotype away from the steeply decreasing side of the fitness function may provide a more useful and realistic case of conservative bet-hedging (CBH). This fulfills the definition of bet-hedging, in that it increases geometric mean fitness at a cost of lower arithmetic mean fitness (i.e. it provides lower fitness in the average environment, but also a lower variance in fitness across environments). Such a type of CBH has not been formally modelled previously, rather theoretical treatments of CBH have been limited to models with two discrete environments, where CBH has been envisioned as a canalized phenotype providing a compromise between the peaks of the fitness functions for the two environments (Starrfelt and Kokko 2012; Crowley et al. 2016). Such a CBH strategy is unsatisfactory since it always loses to a DBH strategy (producing specialists for the two environments) and is therefore not compatible with the concept of a ‘continuum’ between CBH and DBH and thus some sort of coexistence of the two strategies.

Various empirical studies of traits based on skewed fitness functions have invoked CBH arguments of the type we model here (e.g. Boyce and Perrins 1987; Simons and Johnston 2003). However, this shift is often also adaptive from an arithmetic mean fitness point of view in terms of an ‘insurance’ strategy (Dall 2010). With a skewed fitness function, the cliff-edge effect entails that if individuals with the same genotype differ stochastically in their phenotypes (or the fitness value of their phenotypes, e.g. due to inhabiting different microenvironments), their average fitness is maximized if the mean phenotype is shifted away from the peak of the fitness function, towards the less steeply decreasing side (Mountford 1968; Vercken et al. 2012; Mitteroecker et al. 2016). The same is also true for a single individual experiencing uncertainty about its current state (i.e. regarding its phenotype or position on the x-axis on the fitness function) or uncertainty about its current microenvironment (i.e. the position of the fitness function on the x-axis relative to its phenotype). Its average fitness is therefore also maximized by ‘playing it safe’ and moving its phenotype away from the peak down the shallow slope of the skewed fitness function. This shift (insurance) in the mean phenotype thus increases both arithmetic and geometric mean fitness and is not simply a bet-hedging strategy. We initially hypothesized that CBH (lowering the variance in expected fitness for each individual) might shift mean phenotype even more away from the peak of the fitness function than insurance alone. However, our analysis shows that maximizing geometric mean fitness generally does not require a different (i.e. a further shifted) phenotype as compared to the phenotype that maximizes arithmetic mean fitness, but rather there is an exact alignment of fitness interests for short-term and long-term strategies.

An exception to this is in cases where there is suboptimally little phenotypic variance, *σ_k_*. The effect is shown in Fig. 5A, where geometric mean fitness (blue line) is maximized for a higher phenotypic value (the individual fitness function peaks at zero and has its steepest decline for negative values, see Fig. 1) than arithmetic mean fitness (red line). This difference in optimum phenotypic values for canalized traits stems solely from a fitness variance-reducing benefit and can thus be attributed to bet-hedging, and perhaps CBH specifically. However, in this case the diversification bet-hedging (DBH) effect of increasing phenotypic variance instead increases fitness much more than does any possible CBH effect shifting the canalized phenotype to more positive values further down that shallow side of the fitness function. We would therefore expect selection to favor this DBH mechanism to reduce fitness variance (Lande and Arnold 1983), and again no additional CBH is expected to evolve beyond that already captured by the effect of adaptive insurance.

We also conclude that, given the opportunity for insurance, there is very little scope for a single trait to exhibit both DBH and CBH as an additional adaptation on top of any adaptive insurance already being selected for. We do see DBH alongside insurance, but not CBH. An obvious reason for this is that in our model we used the same pattern of environmental stochasticity at both the individual (short-term) and genotypic (long-term) levels. Hence, any adaptive solution at the individual level that maximizes arithmetic mean fitness also accounts for the same regime of environmental stochasticity experienced at the genotype level, leaving nothing left for any geometric mean fitness effects to cope with. This ‘alignment of fitness interests’ maximizing both arithmetic and geometric mean fitnesses in the face of the same regime of environmental fluctuations and skewed fitness functions at the two levels is intriguing, but it does not promise to make empirical evidence for conservative bet-hedging any less ‘elusive’ (Childs et al. 2010; Simons 2011). Whilst the structure of environmental variation in our model might reflect the general pattern expected of environmental stochasticity in nature, regimes of environmental stochasticity may sometimes differ between the within lifetime variation at the individual level versus the across lifetime variation at the genotype level (e.g. seasonal variation versus El Niño events, as experienced by annual organisms). Therefore, understanding adaptations to environmental stochasticity at different levels, such as insurance versus CBH, requires that we appreciate how patterns of the stochasticity in question align and differ at the different levels of organismal experience.

Our result in this regard brings into focus the ecological relevance of previous work. For example, Lof et al. (2012) used a stochastic dynamic model of timing of reproduction in great tits when the timing of the food peak fluctuates between years. They assumed an asymmetric fitness function of laying date relative to the food peak and showed that maximizing expected (arithmetic mean) fitness does indeed produce an adaptive mismatch with the food peak, in the direction away from the steeply decreasing side of the fitness function (an ‘insurance’ result). They acknowledge that “there might be additional benefits of adaptive mismatch in terms of reductions in fitness variance” - i.e. maximizing geometric rather than arithmetic mean fitness might yield a different result if fitness variance is lowest for a different laying date than the observed outcome. However, their forward simulations (using the optimal decision matrix from the dynamic model) “suggest that the variation in fitness often exhibited a minimum close to the observed optimal laying dates” (Lof et al. 2012). Hence, there is little scope for CBH to shift the optimal laying date any further away from the cliff edge. However, geometric mean fitness benefits resulting from a reduction in fitness variance across generations can provide an added selection pressure towards the same optimum phenotypic values as insurance. This result appears in our model in that the geometric mean fitness surfaces are more peaked around the maxima than the arithmetic mean fitness surfaces (Figs 2 & 3) - i.e. the selection pressure toward the same insurance optimum becomes stronger due to also producing lowest fitness variance (i.e. the CBH effect) across generations at this same optimum. All of which may provide reason for optimism with regards to species survival in a period when human-induced environmental change may produce sudden increases in environmental stochasticity that are too rapid for effective evolutionary responses (Barrett and Hendry 2012; Nadeau et al. 2017). This is because in the case of asymmetric fitness functions then any currently adaptive insurance strategy will already have selected for the appropriate phenotype, and no extra evolutionary conservative bet-hedging (CBH) response will be needed in terms of additional changes to the mean phenotype.

The importance of environmental stochasticity at the individual versus genotypic level is highlighted in a large body of previous work on bet-hedging (Levins 1962; Cohen 1966; Gillespie 1974). These identify the ‘grain’ of the environmental variation as a strong determinant of whether bet-hedging strategies will evolve (Starrfelt and Kokko 2012; Crowley et al. 2016). If individuals in a population experience very different environments within a generation (i.e. the environment is ‘fine-grained’ or ‘locally variable’), the fitness correlations between individuals of the same genotype will be low and the scope for DBH is reduced. Assuming a continuous distribution of environmental fluctuations, such as in our model, a very fine-grained environment also implies that a larger proportion of the total environmental variation is experienced by individuals of the same genotype within each generation. The between-generation fluctuations in mean environment therefore become smaller, and a smaller proportion of the variance in fitness is experienced at the genotype level. Only when the environmental conditions are common to a large proportion of the population every generation (i.e. a ‘coarse-grained’ environment, featuring ‘global variation’) is there a large variance in genotype fitness between generations, which can be ameliorated by adaptive bet-hedging strategies.

While environmental ‘grain’ is not explicitly modeled here, it is interesting to consider the *σ_k_* gene as any trait that interacts with the grain of the environment. DBH traits, such as offspring dispersal and variation in dormancy duration (i.e. dispersal in time and space), in effect respond to and modify the grain of the environment as it is experienced. Hence, adaptive dispersal (in time or space) leads such genotypes to experience a more fine-grained environment (Gourbière and Menu 2009; Scheiner 2014). We therefore see why our model suggests that the possibility of DBH makes additional CBH on top of insurance unnecessary − the correlation in fitness between individuals of the same genotype can always evolve to be low enough (via DBH) such that arithmetic mean fitness is a good determinant of long-term fitness.

An early model that hinted at this point involved three distinct adaptations for ‘reducing risk in variable environments’ in seed production in desert plants: seed size, dispersal and dormancy (Venable and Brown 1988). Seed size increases survival in bad environments, but does not affect survival in good environments. Given the trade-off between seed number and size, small seeds are optimal in good environments, but large seeds have a lower variance in expected fitness per individual. Whether increasing seed size represents insurance or CBH depends on the proportion of the environmental variation that is experienced by the genotype within versus between each generation (i.e. the grain of the environment). With no dispersal or dormancy, the environment is coarse-grained, fitness is purely a multiplicative process and increased seed size is clearly a bet-hedging trait increasing geometric mean fitness at the cost of a lower arithmetic mean fitness. But increased seed size may also maximize arithmetic mean fitness across fine-grained environments, which is the appropriate fitness measure if the genotype is sufficiently spread in space (or time, in this case) to experience the full range of environmental variation in each generation (Levins 1962). Thus, dispersal and dormancy are not only diversifying bet-hedging (DBH) traits, but also determine the grain of the environment and the need for conservative bet-hedging (CBH) versus insurance. Venable and Brown (1988) show that these adaptations to reduce risk are essentially substitutable and that a decrease in the value of either of the traits away from the optimum leads to evolutionary compensation in an increased value of the other traits. However, note that the DBH traits here, dispersal and dormancy, are inherently different from a conceivable fourth risk-reducing strategy: variation in seed size itself. In discrete environments (‘good’ and ‘bad’) the optimal DBH strategy maximizing geometric mean fitness is to produce seeds with optimal size for each of the environments (small and large) with the probabilities of those respective environments occurring, while in continuously varying environment seed size should vary around a mean trait value, as in our model. Here we showed that in a continuously varying environment, this type of DBH (which Venable and Brown (1988) do not explore) generally provides a greater benefit than CBH. However, we do not rule out that with other types of environmental variation and environment-specific fitness functions then changing the mean trait value may be a better strategy. We also point out again that both of these risk-reducing strategies are only effective once the grain of the environment causes selection to maximize geometric rather than arithmetic mean fitness (Venable and Brown 1988; Scheiner 2014).

In our current model, the only time arithmetic mean fitness would be higher with more phenotypic variance (i.e. for an individual, rather than a DBH genotype increasing among-individual variation) would be if its fitness function were strongly convex around its current phenotype (Fig. 5b), such as at the tails of a Gaussian fitness function. This is the same adaptive gambling effect that produces risk-sensitivity (Caraco et al. 1980; Stephens 1981), aka variance sensitivity (Smallwood 1996; Stephens et al. 2007), which is an important concept from economics used to explain foraging decisions and other behaviors when there are more or less variable options. The fitness advantage of variance sensitivity follows directly from Jensen’s inequality: if the fitness function *f* of some utilized resource or trait *x* is convex, then the mean fitness gained over a sequence of events with variable reward *x* will be larger than the fitness gained from the mean reward *x*: E[*f*(*x*)] > *f*(E[*x*]). This is an arithmetic mean fitness maximizing argument, and the benefit of increasing phenotypic variance can therefore be seen in our calculations of arithmetic mean fitness as well (top rows of Fig. 2 & 3; Fig. 5b). Essentially, for a constant *μ_k_* sufficiently far from the fitness function peak, arithmetic mean fitness is maximized at an intermediate value *σ_k_* > 0. This similarity between variance-sensitivity and DBH in producing variable phenotypes but at different adaptive timescales has not been reported before, and it is made explicit here through our comparison of trait means and variances maximizing long-term arithmetic or geometric mean fitness.

We have demonstrated several results linking theory concerning individual-level strategies from behavioral ecology with genotype-level adaptations from evolutionary biology in context of environmental uncertainty. The next step is now to connect these theoretical studies to real world examples and quantitative studies of organisms in the lab and in the wild, especially if we are to understand how populations might respond to current human-induced rapid environmental change. In order to apply our genotype-level view to predictive statements concerning evolutionary responses we would require extensive data on past climate fluctuations, clear links between trait values and individual fitness, as well as detailed knowledge of the genetic mechanisms underlying the traits. While this might seem an insurmountable task, the empirical evidence for bet-hedging in the wild has shown that long-term studies on natural populations can provide answers to these types of questions (Simons 2011). We hope that our results here act as a motivation to both empirical and theoretical studies on adaptations to stochastic environments to compare and contrast individual versus genotype perspectives and the alternative adaptive currencies of arithmetic versus geometric mean fitness.

## Acknowledgements

This work was supported by the Research Council of Norway through its Young Talented Researchers funding scheme FRIMEDBIO, project number 240008 to I.I.R., and partly through its Centres of Excellence funding scheme, project number 223257, to Centre for Biodiversity Dynamics at the Norwegian University of Science and Technology.

## Author contributions

IIR, JW and JT conceived of the ideas and all authors contributed to the development of the ideas. JT and TRH wrote the code and did the calculations. TRH wrote the manuscript with contributions from all authors. All authors gave final approval for publication.

## Code accessibility

The code is available upon request, and will be uploaded to Dryad upon publication.

